# Detection of Autism Spectrum Disorder Using Graph Representation Learning Algorithms and Deep Neural Network, Based on fMRI Signals

**DOI:** 10.1101/2022.06.23.497324

**Authors:** Ali Yousedian, Farzaneh Shayegh, Zeinab Maleki

## Abstract

In this paper, we are going to apply graph representation learning algorithms to identify autism spectrum disorder (ASD) patients within a large brain imaging dataset. Since ASD is characterized by social deficits and repetitive behavioral symptoms, it is mainly identified by brain functional connectivity patterns. Attempts to unveil the neural patterns that emerged from ASD are the essence of ASD classification. We claim that considering the connectivity patterns of the brain can be appropriately executed by graph representation learning methods. These methods can capture the whole structure of the brain, both local and global properties. The investigation is done for the brain imaging worldwide multi-site database known as ABIDE (Autism Brain Imaging Data Exchange). The classifier adapted to the features embedded in graphs is a LeNet deep neural network. Among different graph representation techniques, we used AWE, Node2vec, Struct2vec, multi node2vec, and Graph2Img. The best approach was Graph2Img, in which after extracting the feature vectors representative of the brain nodes, the PCA algorithm is applied to the matrix of feature vectors. Although we could not outperform the previous 70% accuracy of 10-fold cross-validation in the identification of ASD versus control patients in the dataset, for leave-one-site-out cross-validation, we could obtain better results (our accuracy: 80%). It is evident that the effect of graph embedding methods is making the connectivity matrix more suitable for applying to a deep network.

## 1. Introduction

Autism spectrum disorder (ASD), is a set of clinical presentations, emerging due to neurodevelopmental disorder. ASD symptoms are related to social communication, imagination, and behavior. Early diagnosis and early intervention before age 4 (e.g., 12–48 months) can improve cognition, language, and adaptive behavior significantly (Elder et al., 2017). Accurate and timely diagnoses of ASD significantly improve the quality of life of individuals with ASD. Although there has been widespread research in this field, there is no clear etiology to diagnose ASD.

To date, ASD diagnosis is done based on the behavioral characteristics of children, observed by parents and teachers at home or school (Nickel & Huang-Storms, 2017) (Almuqhim & Saeed, 2021). Since autism is related to abnormal development of the brain, assessing the brain function is at the top of automatic diagnosis and classification research. Resting-state functional MRI has been suitable, due to its spatial resolution. Brain regions interact together to emerge an especial behavior, *i.e*., a region’s function is tightly dependent on its interactions. Various differential observations consider the properties of the brain network in healthy and ASD subjects. However, statistical analysis could lead the researchers to a disorder whose mechanisms vary among patients. In other words, there is no unique fact to announce them as biomarkers of ASD.

By considering brain regions and their connections as a network, autism spectrum disorder detection alternatively could be a network classification task, in which machine learning techniques could help. To efficiently use information hidden in the resting-state fMRI, the connectivity measures obtained from resting-state fMRI are useful to understand the large-scale functional difference between healthy and abnormal brains.

After that, a suitable classifier should be used. Vast number of mental disorder diagnosis studies used traditional classifiers, such as Support Vector Machine (SVM), LASSO, and Bayesian classifier. But, deep learning methods showed a major preference in the case of connectivity matrix, because it is a high-dimensional feature of brain activity. The reason is that, a combination of different functional connectivity metrics increases number of hyperparameters of a machine learning algorithm, in such a way that just deep neural networks can afford it, and learn more complex structures of data. From this point of view, applying fully-connected deep neural networks, and convolutional networks on fMRI volumes and raw connectome data appears to be successful (Heinsfeld et al., 2018) (Sherkatghanad et al., 2020).

In (Heinsfeld et al., 2018) deep learning algorithm using the full connectivity matrix is applied to classify ASD and controls using ABIDE data. They showed anterior-posterior underconnectivity in the autistic brain and surpassed the state-of-the-art classification of autism by achieving 70% accuracy. Also, Convolutional Neural Network (CNN) was used to effectively diagnose Alzheimer’s disease (AD) (Sarraf et al., 2016) and mild cognitive impairment (MCI) (Meszlényi et al., 2017). In another study, CNN was used to extract features from fMRI data, and SVM was used for classification (Nie et al., 2016). A Deep Auto Encoder was used to classify fMRI data of MCI (Suk et al., 2016). Furthermore, different hidden layers in between the encoder and the decoder (Patel et al., 2016), were added to afford different tasks, like denoising (Heinsfeld et al., 2018), or generating sparse features (Guo et al., 2017). Other networks, like Radial Basis Function network (RBFN) (Vigneshwaran et al., 2015), Restricted Boltzmann Machine (RBM) (Huang et al., 2016), and Deep Boltzmann Machine (DBM) can be used to extract features from fMRI data, because they can combine the information of different voxels of the region of interest (Zafar et al., 2017).

Different traditional and deep machine learning algorithms reported good performances in ASD classification of ABIDE dataset while considering individual sites (Sherkatghanad et al., 2020). But our main concern is that after intermingling all the sites, or leave-one-site-out cross-validation algorithm, accuracy, *i.e*., the percent of correctly classified subjects, and the area under ROC is diminished. In other words, yet there is not an algorithm appropriate for clinical usage. In summary, the accuracy of some ASD classifiers using SVM and DNN algorithms, some decomposition techniques like Principal Component Analysis (PCA), and graph neural network ranges from 55% to 70.4% (Kazeminejad & Sotero, 2020; Parisot et al., 2018; Sharif & Khan, 2021; Xing et al., 2019). Thus, still, further experiments are required to be conducted with patients of different ages to ensure the clinical value of these methods (Li et al., 2018).

On the other hand, the benefit of DNN is mainly due to a large number of training examples (Guo et al., 2017; Heinsfeld et al., 2018; Kim et al., 2016; Kuang et al., 2014). Developing deep learning approaches to work with functional connectivity (FC) features using small or at best modest sample sizes of neurological data (A. di Martino et al., 2014; Guo et al., 2017; Heinsfeld et al., 2018; Kim et al., 2016; Kuang et al., 2014) is debatable from the reproducibility and generalizability point of view. One solution is the deep transfer learning neural network (DTL-NN) approach that could achieve improved performance in classification for neurological conditions (70.4% for ASD detection), especially where there are not large neuroimaging datasets available (Li et al., 2018). Other solutions are Synthetic Minority Oversampling Technique (SMOTE) to perform data augmentation to generate artificial data and avoid overfitting (Eslami & Saeed, 2019), and sparse autoencoder (SAENet) that was used for classifying patients with ASD from typical control subjects using fMRI data (70.8% accuracy, and 79.1% specificity) for the whole dataset as compared to other methods (Almuqhim & Saeed, 2021). Another approach is to develop a machine learning approach with a robust training methodology (Li et al., 2018). Machine learning algorithms able to extract replicable, and robust neural patterns from brain imaging data of ASD patients, reach suitable classification results (Pereira et al., 2009).

Another solution in studies with limited sample sizes is the reduction of the size of features indicating useful connectivity properties by network analysis methods. The ease of representing brain connectivity information according to graph theory makes them very valuable tools in this area. Machine learning on graphs finds its importance here: Finding a way to represent, or encode graph structure is the subject of this task. Nowadays, in order to model information underlying the graph structure, there are new ways of representing and analyzing graphs, which afford the complexity of working with big graphs. Referring to these representation algorithms as embedding, applying these approaches to brain networks is named connectome embeddings (CE). These embedding algorithms involve converting graphs into vectors. Network embedding techniques can be divided into three buckets: 1) based on engineered graph features, 2) obtained by training on graph data, 3) obtained by a layer of a deep network. The main drawback of the former is that structural homologies or higher-order relations of the connectivity matrix could not be captured (Rosenthal et al., 2018). Furthermore, these features are not flexible, *i.e*., they cannot adapt during the learning procedure. In summary, many of these local and global features cannot capture the topological shape of the graph.

In the second bucket, referred to as shallow embedding, network embedding vectors are learned by optimizing different types of objective functions defined as a mapping to reflect geometric information of graph data. This optimum embedded space is the final feature vector. These algorithms involve learning approaches that map nodes to an embedding space. Anonymous walk Embedding (AWE), Node2vec, Struct2vec, Deep Walk, multi node2vec, and graph2image (Grover & Leskovec, 2016; Ribeiro et al., 2017) are some well-known algorithms of this bucket. Representing higher-order features of the connections of a graph is referred to as the Graph2image method, in which the embedded space of the brain network is transformed into an image. This method is helpful to develop an image that is convenient for training a CNN. The advantage of this method is the capability of dimensionality reduction of this image by an algorithm like PCA, and still have an image at hand (Meng & Xiang, 2018). Multi-node2vec was applied on fMRI scans over a group of 74 healthy individuals. Multi-node2vec identifies nodal characteristics that are closely associated with the functional organization of the brain (Wilson et al., 2018).

In the third bucket, referred to as deep embedding, CE and deep learning algorithms are combined to form a single deep network. This combinatory network can exploit the connectome topology. In this category, a Hypergraph U-Net (HUNet), Graph U-Net (GUNet) (Gao & Ji, 2019) and Hypergraph Neural Network (HGNN) (Feng et al., 2019) are proposed in which low-dimensional embeddings of data samples are learnt from the high-dimensional connectivity matrix. Indeed, these networks emerged as a subset of deep graph neural networks (GNN) (Gao & Ji, 2019; Kipf & Welling, 2017; D. Wang et al., 2016) are able to model the deeply nonlinear relationship node connectomic features (Banka & Rekik, 2019; Bessadok et al., 2019; Ktena et al., 2017).

In this paper, we demonstrate the role of the second bucket (CE method) in representing the structure with which brain regions are connected to each other and assess its effect in ASD classification. Based on the ABIDEI and ABIDEII public datasets, recorded at some different sites, we want to investigate whether CE can surpass previous researches or not. Accordingly, by using CNN classifiers, we claim that there is great potential in combining graph representation methods, with deep learning techniques for fMRI-based classification.

In fact, we claim that representation-based features can solve the problem of high-dimensional input of the deep network. The task of classification based on a connectivity matrix is very similar to image classification. In this matrix, each row and column correspond to a brain region. The value of every entry is the connectivity measure between two relevant regions which its row and column stand for. However, the matrix obtained by connectivity embedded patterns is not exactly similar to natural images. In this matrix, any shift of the patterns is interpretable. Thus, the design of the convolutional filters is different from those of natural images, because they should consider the properties of the connectome database to determine different connectivity structures. Therefore, our idea to use the information of node structures as a new and low-dimensional source might increase classification performance. However, such dimension reduction may lead to more ambiguity about the place of alteration in the connectivity matrix.

The structure of the paper is as follows: After describing the network embedding techniques in section 2, suitable embedding-based features are illustrated. In section 3, classification technique using deep network is declared. Afterward, ABIDE database, their pre-processing methods, and the embedded features extracted from them are introduced. These features are applied to deep network to detect ASD subjects. Some evaluation measures like F-score, and accuracy of this classifier is reported in the Results section and is compared to other literature working on ABIDE dataset.

## 2. Network Embedding Concept

The concept of network embedding date back to the skip-gram model, originally proposed for word embedding. It can be described as follows: Suppose there is a graph *G* = (*V, E, A*) with *V* as the node set, *E* as the undirected and weighted edge set, and *A* as the adjacency matrix. We are going to find the optimum function *z* = *f*(*ν*) ϵ *R^d^* that maps each node or sub-graph to a d-dimensional vector disclosing the structure of the graph. These vectors should be representative of the graph and can be used as the feature vectors uncovering the similarities of the graph for machine learning algorithms. At this level, each node corresponds to a d-dimensional embedded vector involving its connections with all other nodes (Hamilton William L. et al., 2017).

Indeed, these low-dimensional embedded vectors can summarize either position of nodes in the graph, or the structure of their neighborhood, and user-specified graph statistics (Hamilton William L. et al., 2017). Most shallow embedding mapping techniques are done based on a look-up table, just like what is occurred in classic matrix factorization for dimensionality reduction (Hamilton William L. et al., 2017). For another part of shallow embedding techniques, learning the embedded vector for each node is in fact training an encoder-decoder system, defined as an optimization method. The decoder maps the similarity of two nodes into a real-valued similarity measure. Different techniques able to afford this job (like Deepwalk, Node2vec, AWE, TSNE, GraRep, and others) are based on a stream of randomly generated walks. Random walk methods lead to better performance, due to their flexible and stochastic nature of the similarity they consider. The resultant vectors can describe the similarities and subgraph membership with relatively few dimensions. These learned embedded vectors can be used as features of the graph.

The core of this relevant optimization problem is to find a mapping such that nearby nodes in short random walks have similar embedding vectors. As a similarity measure, *P_g_*(*ν_j_*|*ν_i_*) is the probability of visiting *ν_j_* on a random walk starting from *ν_i_* and with length *T*. For each vertex *ν_i_*, a sequence of vertices is formed, from which the probabilities can be obtained, according to the ratio of weight of edge (*ν_i_*,*ν_j_*) to the total (positive) edge weight in the *r*-truncated random walks. Here *r* varies from 1 till the original walk length *T*. As seen in Fig. 1, nearby nodes are mapped into vectors with small between angles.

**Figure 1.**
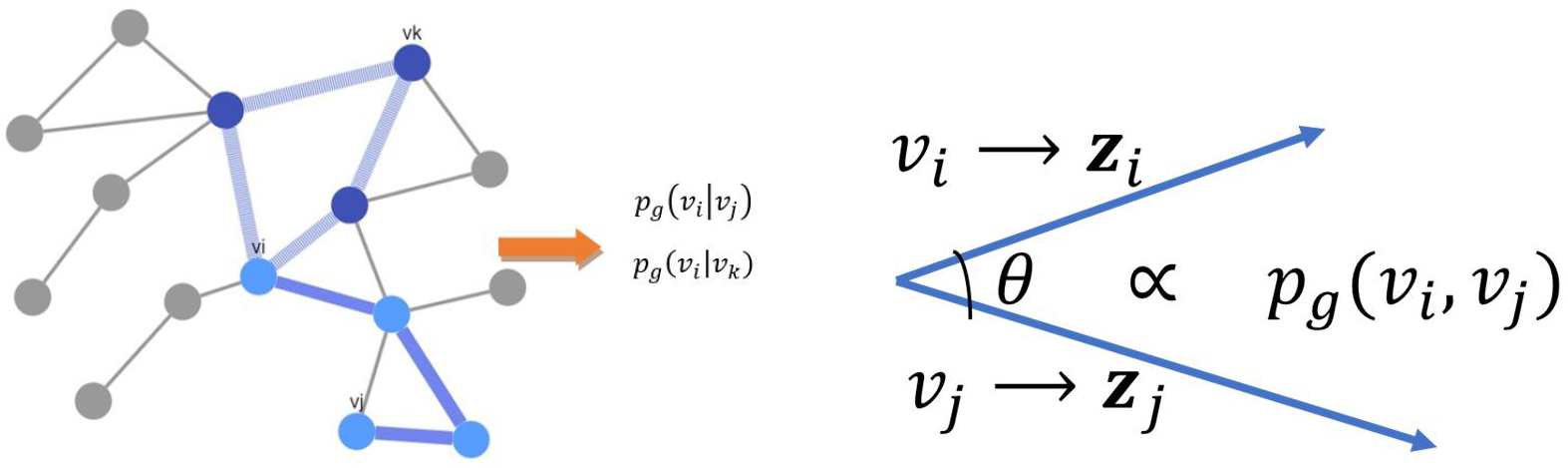
Schematic of random-walk as the basis of shallow embedding techniques. Possible random walks between every two nodes lead to obtaining the conditional probability of these nodes. This probability is converted into the angle between the embedded vectors of these nodes (Hamilton William L. et al., 2017)

### 2.1 Node2vec and DeepWalk

Two of the well-known network embedding algorithms defined based on the optimization methods are Node2vec and DeepWalk. For DeepWalk and Node2vec algorithms, decoding is done to obtain vectors *z_i_, z_j_* according to an algorithm defined as below (Grover & Leskovec, 2016):

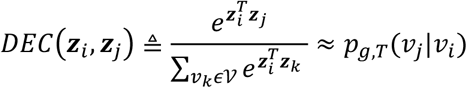

where *P_g,T_*(*ν_j_*|*ν_i_*) is the probability of visiting *ν_j_* on a random walk starting from *ν_i_* and with length *T*. Both Node2vec and DeepWalk algorithms could be easily described by the concept of context graph. This context graph is an auxiliary graph *C* in which the weight of edge between the source node *ν_i_*, and its context *ν_j_*, *i.e*., *c_i,j_* is proportional to the similarity between nodes *ν_i_* and *ν_j_* in graph *G*. No edge between them, is equivalent with weight −1. For a directed graph *G*, determines the role of a vertex as sink or source node. Based on context graph and optimization, *P_g,T_*(*ν_j_*|*ν_i_*) is determined. However, there are different network representation learning algorithm. But, among them Node2vec and DeepWalk are the most popular ones (Khosla et al., 2021).

In Node2Vec and DeepWalk algorithms similarity of nodes *P_g,T_*(*ν_j_*|*ν_i_*) is obtained via assessing the nodes sampled in random walks through graph. In other words, weights of edges of context graph *C* is the number of walks able to reach node *ν_i_* via *ν_j_*. Sampling of the nodes on which random walk move, is different in Deep-walk and Node2vec (Khosla et al., 2021). While DeepWalk performs a uniform random walk, Node2vec follows a 2^nd^ order biased random walk. In other words, in Node2vec algorithm, negative sampling (Z. Yang et al., 2020) is used, but in DeepWalk algorithm, hierarchical softmax technique is employed. Distribution function underlying the probability of visiting a node in the random walk differ in these two techniques. Thus, the optimization cost function defined as an expectation of all combinations of elements, and subsequently the results, differs in these two algorithms.

In Fig. 2, bread-first and depth-first search (BFS and DFS) in a graph for node *ν*\* (the algorithms used in random walk) are shown. In BFS the neighborhood is restricted to immediate neighbors of *ν*\*, *i.e*., for a neighborhood of size *k* = 3 the nodes *ν*_2_, and *ν*_3_. In DFS, the neighborhood consists of nodes sequentially sampled at increasing distances from *ν*\*. In Fig. 2, DFS samples *ν*_4_, *ν*_5_, *and ν*_9_.

**Figure 2.**
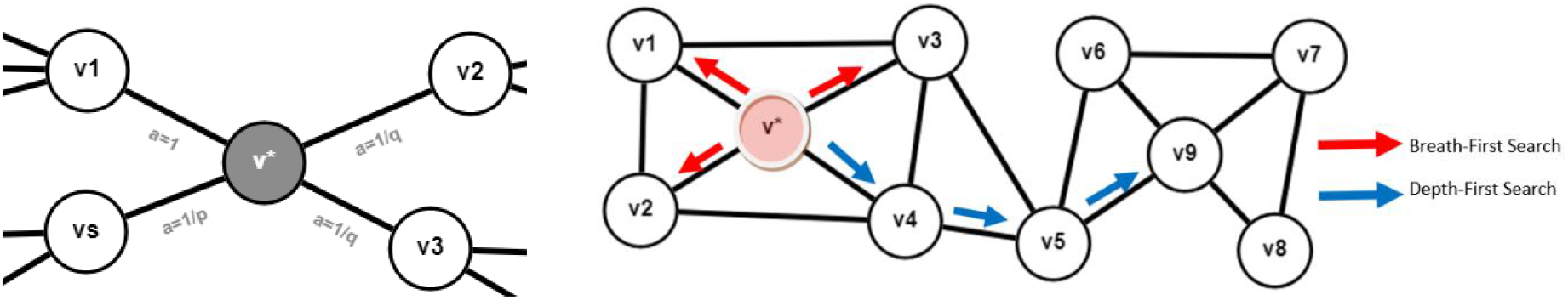
A. The way Node2vec bias the random walk by p and q, B. Difference between Breath-First Search (BFS) that find one-hop neighborhood, and Depth-First Search (DFS) random walks that explore further away, to capture the structure.

Node2vec does an interpolation between BFS and DFS via two parameters *p* & *q*, selected suitably. Hyperparameters (*p* & *q*) control likelihood of walks immediately revisiting a node, and revisiting a node’s one-hop neighborhood, respectively (Khosla et al., 2021). In a biased random walk, the probability of transition from a node to the next one is different according to the length of the shortest path between them (Grover & Leskovec, 2016). This can be generalized to higher order representation by considering the sequences would be appeared during random walks gathered in set *D*, including triples of nodes, with parameters *p* & *q* (Khosla et al., 2021). Thus, probability of traveling from node *u* to *v* and then *w* indicated by *<u>P</u>_u,v,w_* can be defined as below.

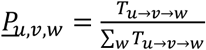

Where

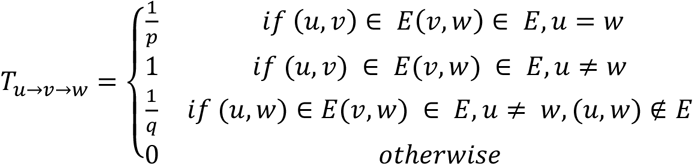

where *E*(*v, w*) is the shortest path between *v* and *w*. This algorithm is summarized in Algorithm 1. As well, DeepWalk algorithm is illustrated in Algorithm 2.

Node2vec and Deep-walk approaches lead to a unique embedding vector for every individual node, but have some drawbacks, including woring as a lookup table, its computational cost, fail to leverage attribute information of nodes involving node’s position and role, weakness in predict information of unseen nodes. To alleviate the abovementioned drawbacks, two alternatives are arisen: 1) some embedding approaches that enable capturing the structural roles of nodes, have been proposed (Donnat et al., 2018; Ribeiro et al., 2017), 2) Network embedding in a feature-based manner has been proposed.

#### Algorithm 1 Node2Vec Algorithm (Meng & Xiang, 2018) (Grover & Leskovec, 2016)

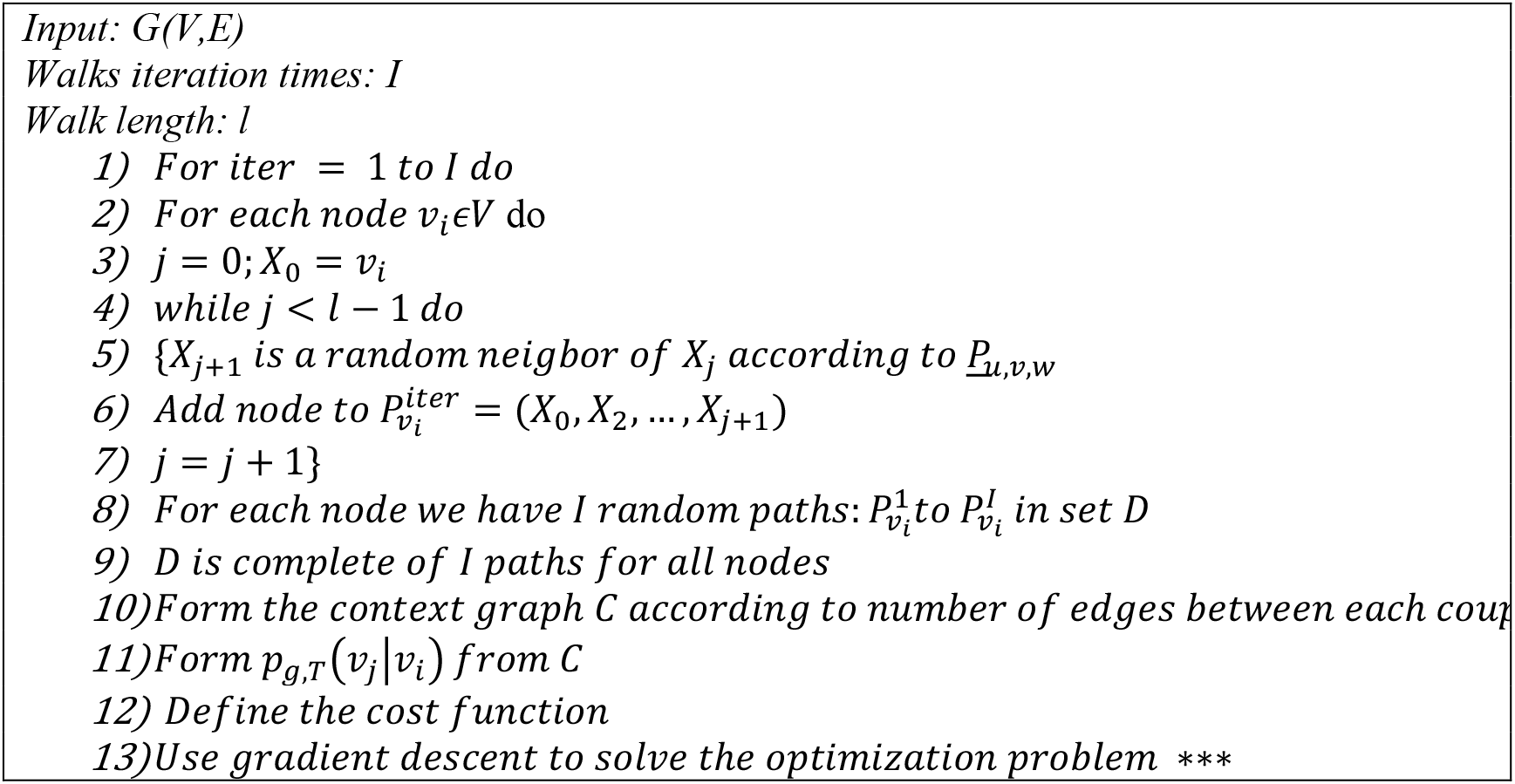

#### Algorithm 2 DeepWalk Algorithm (Perozzi et al., 2014)

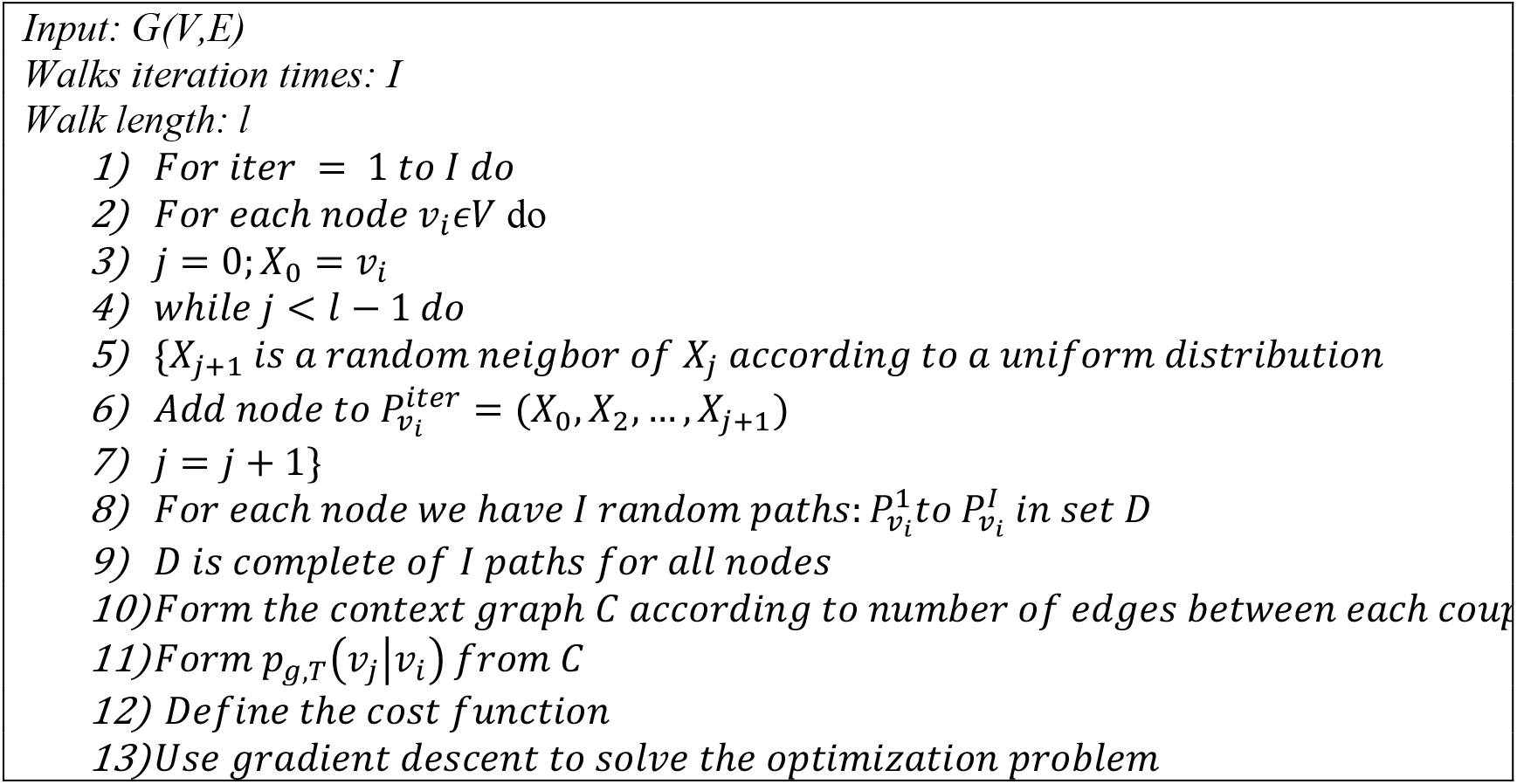

### 2.2 Struct2Vec

As an example of the first alternative, Ribeiro technique referred to as Struc2Vec, generate some new graphs *G*_0_, …, *G_K_*, each to capture one kind of *k*-hop neighborhood structural similarity, from the original graph *G*. The algorithm is as follows (Ribeiro et al., 2017):

1. For each node *ν_i_*, order the sequence of degrees of nodes exactly with distance of *k*-hops from it: *R_k_*(*v_i_*).
2. Start from a weighted graph *G*_0_ whose edges have zero weights *w*_0_(*v_i_, v_j_*) = 0.
3. Build a sequence of weighted graphs whose edges varies adaptively by the equation:

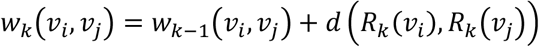

where *d*(*R_k_*(*ν_i_*), *R_k_*(*ν_j_*)) is distance between sequence *R_k_*(*ν_i_*) and *R_k_*(*ν_j_*), and could be defined with different measures.
4. Run random walk on new graphs *G*_0_,…, *G_K_* to implement Node2vec on them, and learn latents from them, using an algorithm like SkipGram.

### 2.3 Feature-based embedding

In the second alternative, namely, feature-based methods, two algorithms referred to as Graph2Img and AWE are considered.

The Graph2Img algorithm, first transfers the original network into feature vectors and then uses clustering methods to group nodes. In other words, after graph node embedding process into a *d*-dimensional space, representations of nodes are gathered in a matrix of dimension *N*×*d*, where *N* = |*V*|, *i.e*., number of nodes in the graph. Next, we can decide whether all features are important or not, and determine their priority. In fact, we can use just the most important dimension, the second important one, and so on. Here, principal component analysis (PCA) method is used to transform the *d*-dimension vector of node *ν_i_*, into dPCA-dimension vector LPCA, which is a sequential list.

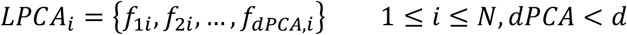

where *N* represents the number of nodes in the graph. In such a way, for each component *f*_1_ there are a *N*-dimension vector. Thus, for each pair of components, we can build a matrix of 2 × *N* dimension, and interpret it as *N* ordered pairs, or *N* points in a 2D plane. As an example, *M*_12_ is built from components 1 and 2:

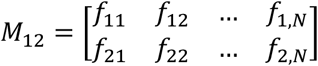

Digitizing values of LPCA components to *r* equally sized bins, this two-dimensional space is partitioned into *r* × *r* bins, i.e., there are *r* × *r* potential states for an ordered pair. 2D histogram of these ordered pairs (namely *H*), is a *r* × *r* matrix, in which *H_ij_* denotes the count of node falling into (*i,j*) state.

#### Algorithm 3. Graph2Imag Algorithm

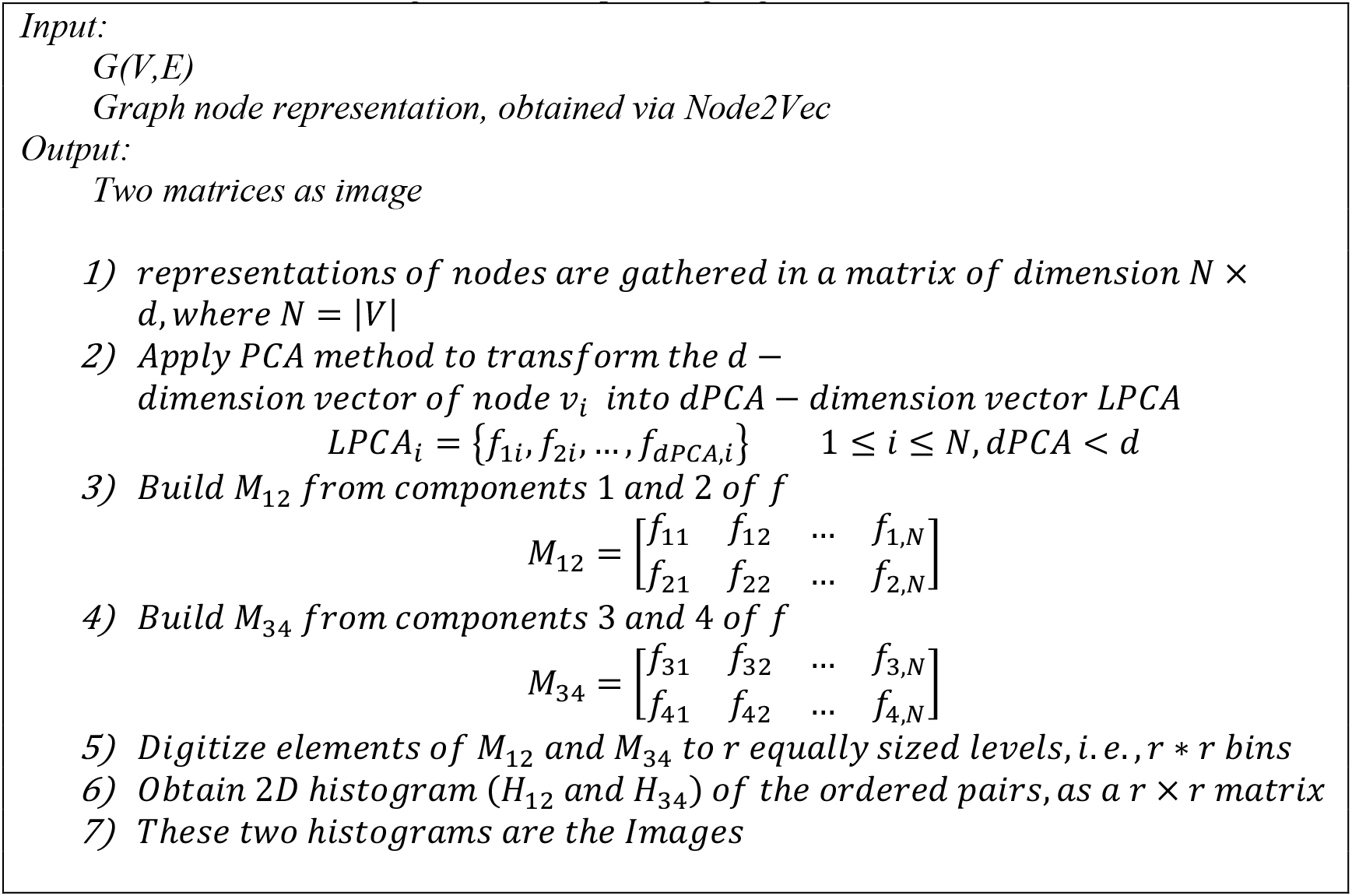

Among all dPCA*dPCA matrices *M*, could be built by this procedure, we just consider *M*_12_ and *M*_34_, *i.e*., taking into account just four first components of LPCA, which seems to be enough to analyze the brain network. As shown in Fig. 3 (Meng & Xiang, 2018) these two matrices, behaving like images, can be applied as different channels of DCNN. The algorithm pseudo-code is shown in Algorithm 3 (Meng & Xiang, 2018).

**Figure 3.**
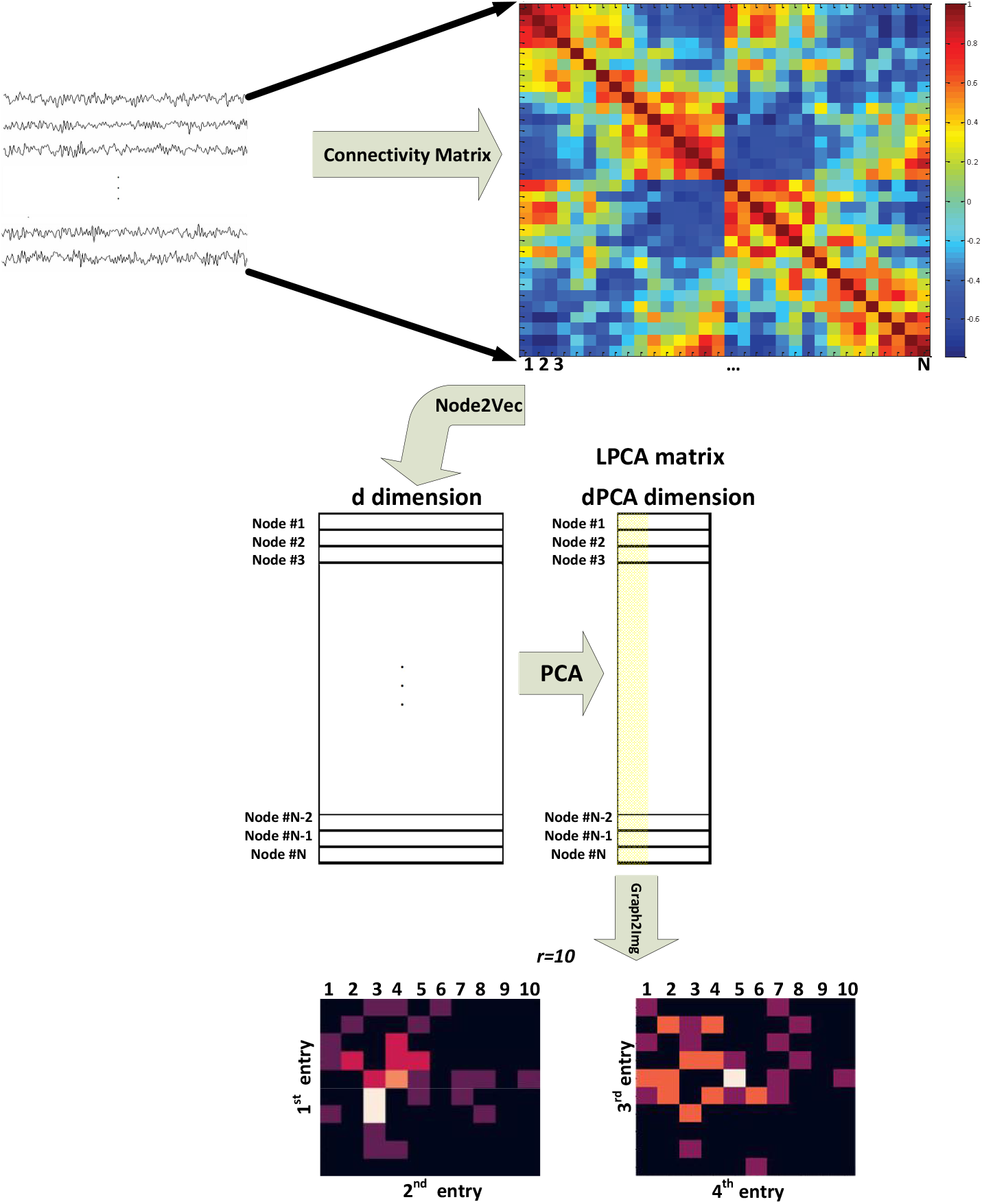
block diagram of the Graph2Img Feature-based embedding algorithm (Meng & Xiang, 2018)

As another feature-based network embedding method, Anonymous walk Embeding (AWE) algorithm used distribution of anonymous walks. Anonymous walks are the set of walks starting from an initial node *u*, by length *l* passing from random nodes, and termination at node *ν*. There are a set of *η* such random walks 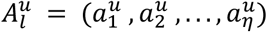. Thus, number of all possible random walks with length *l* exponentially grows with *l*. These anonymous walks capture structural information of nodes, because labels of the nodes constituent of a random walk is omitted for them. In fact, corresponded to the random walk: *w* = (*ν*_1_, *ν*_2_, …, *ν_k_*), we can define an anonymous walk involving sequence of integers *a* = (*f*(*ν*_1_), *f*(*ν*_2_),…, *f*(*ν_k_*)) where *f*(*ν*) is the minimum place of *ν* in the *w* random walk (Ivanov & Burnaev, 2018). However, due to huge number of anonymous walks of a large graph, an efficient sampling approach is required to approximate this distribution (Ivanov & Burnaev, 2018). Defining the objective function of similar nodes on local neighborhoods of anonymous walks, improve the structural consideration of the embedding method.

AWE is defined on weighted directed graphs. Given a weighted graph *G* = (*V, E,A*), a random walk graph could be constructed *R* = (*V, E, P*) in which every edge *e* = (*u, v*) has a weight *p_e_* (Ivanov & Burnaev, 2018). A random pair of nodes (*u_i_*,*u*_*i*+1_) can be led to a random walk with probability of *p_e_*(*u_i_,u*_*i*+1_). Accordingly, it is possible to attribute *p*(*w*) to each random walk; as the multiplication of probability of choosing total pairs in that walk, *i.e*., 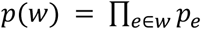.

Starting from each node, we can obtain marginal probability of selecting a special walk according to *p*(*w*). A probability of seeing anonymous walk 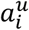 of length *l* for a node *u* is 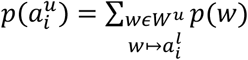 where *W^u^* is the set of all walks starting from *u*, and 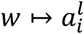 involves the walks with length *l*. Anonymous walk embedding of this set of all possible anonymous walks of length *l* of graph *G* is a vector whose *i*th component is the probability of that walk. Aggregating probabilities across all vertices in a graph and normalizing them by the total number of nodes N, we get the probability of choosing anonymous walk *a_i_* in graph *G*:

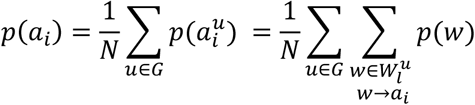

AWE algorithm includes computation the probability of all random walks (Ivanov & Burnaev, 2018). Since direct computation of AWE requires determining all different random walks in graph *G*, grow exponentially with the number of steps *l*, the AWE algorithm is as follows (Ivanov & Burnaev, 2018):

1. Beginning from each node *u*, sample *N* random walks starting from *u*, and map it to an anonymous random walk.
2. Gather all these sampled anonymous random walks for all nodes. These are collection of co-occurring walks.
3. Learn representation vector of these walks, and generate a *η* * *da* dimensional matrix (*W*), where *η* is the number of all possible anonymous walks of length *l*, and *da* is the embedding size. This model is used to predict a target vector for anonymous walks in a graph.
4. For a graph vector *d*, according to sampled co-occurred anonymous walks in *W*, the model calculates a probability function to predict a target walk among all sampled anonymous walks
5. Now model updates the matrix *W* and graph vector *d* via gradient backpropagation, as schematically is shown in Fig. 4.
6. After repeating steps 4 and 5, a learned graph vector d is called anonymous walk embedding.

**Figure 4.**
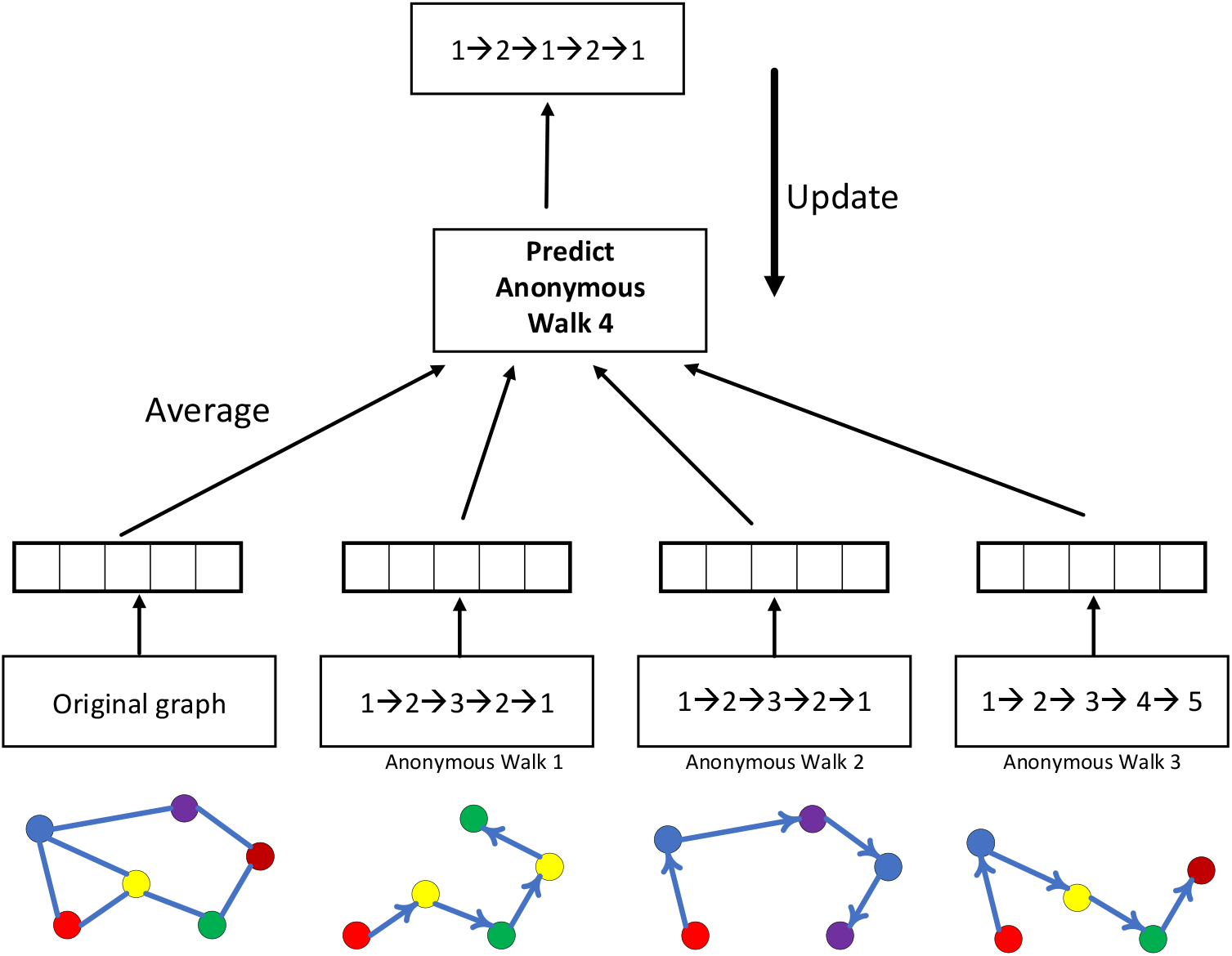
Schematic illustration of learning structure via anonymous random walks (Ivanov & Burnaev, 2018)

These four embedding algorithms, Node2Vec, DeepWalk, AWE, and Graph2Img, extract the feature vectors of each node, describing the characteristics and structure of the graph. Thus, the next step of our research is classification of these feature vectors obtained for healthy and ASD subjects.

## 3. Classification

Graph classification is a task to predict whether a whole graph belong to any class of *C* pre-defined classes. In other words, the task is to train a classifier based on *N* graphs {*G_i_*}, *i* = 1:*N* and their corresponding labels {*L_i_*}, *i* = 1: *N*, able to classify every new graph *G* → *L*. Graph classification problem can be done using two typical approaches: (1) classification using extended CNNs to be appropriate for the raw graphs (Niepert et al., 2016) and (2) graph kernel methods (Shervashidze et al., 2011), in which graph embeddings *f*(*G*_1_) are used in conjunction with kernel methods (*K*(*f*(*G*_1_), *f*(*G*_2_))) to perform classification of new graphs, where *K*: (*x, y*) → *R^n^* is a kernel function, quantifying distance of graphs.

As mentioned earlier, the aim of this paper is kernelized classification of healthy and autistic patients based on functional connectivity matrices. The features extracted from these matrices (*f*(*G*_1_)) are the embedded vectors obtained by using Node2vec, Struct2vec, AWE, and Graph2Img algorithms. To do classification job, we used the DNN classifier. The reason underlying this selection is size of the resultant feature vectors, whose classification requires many parameters to be trained. As well, to validate the performance of our classification task, cross-validation is applied.

Three types of deep network have been considered in this study: LeNet, ResNet, and VGG16. However, finally, we have used LeNet, because of its best performance for our problem. Thus, we just describe it here.

### 3.1 LeNet

*LeNet*, one of the first published CNNs in computer vision tasks, was introduced by (and named for) Yann LeCun. In 1989, LeCun published the first study in which he could train CNNs via backpropagation. Then this network was applied in AT&T Bell Labs, for the purpose of recognizing handwritten digits in images (LeCun et al., 1998). LeNet achieved outstanding results comparable with that of support vector machines, and thus became a dominant approach in supervised learning.

LeNet (LeNet-5) consists of two parts (G. Wang & Gong, 2019): *(i)* a convolutional encoder, and *(ii)* a dense block. The former consists of two convolutional blocks, and the latter consists of three fully-connected layers; The architecture is summarized in Fig. 5.

**Figure 5.**
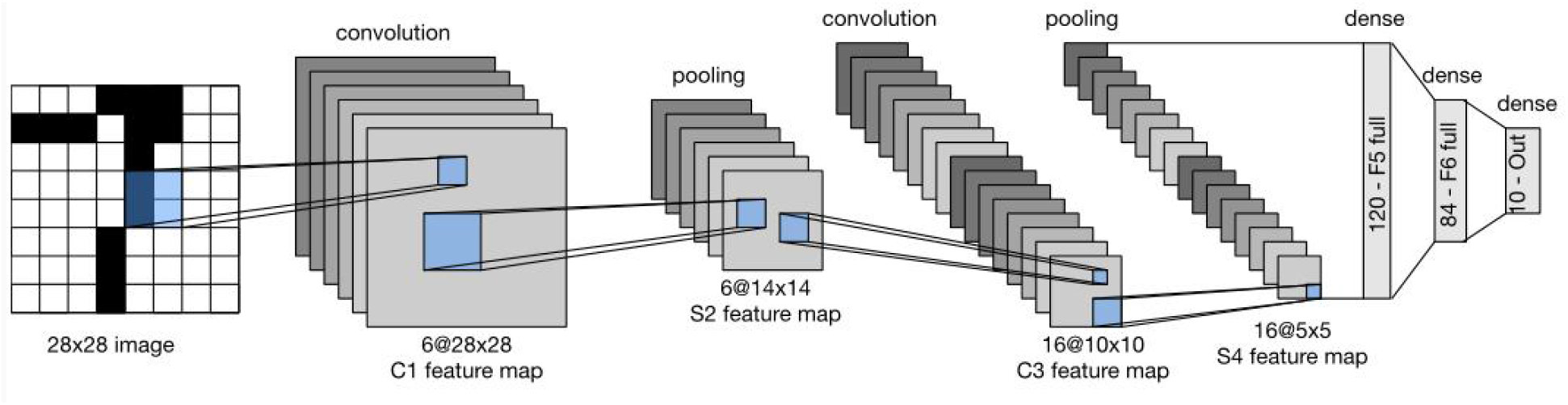
Data flow in LeNet. The input is an image, the output a probability over different possible outcomes (Loey et al., 2016).

Each convolutional block includes a convolutional layer, a sigmoid activation function, and a subsequent average pooling operation. In 1990 ReLUs and max-pooling were discovered to have suitable performance. However, in LeNet, each convolutional layer maps any 5×5 part of the input to a scalar using a kernel and a sigmoid activation function. There is 6 convolutional layers, in such a way that the result is a 6@28*28 tensor. In fact, by these convolutional layers, spatial features of input are mapped to a number of two-dimensional feature maps, namely, channels. Then, a pooling layer sample the channels by a factor of 2, and lead to a 6@14*14 array. Then there is another convolutional layer. Again, this is a convolutional layer with 5*5-dimensional filter. First convolutional layer had 6 output channels, while this second layer has 16 outputs of size 10*10. The output of the convolutional block must be flattened before passed to the dense block. This output is a 16@5*5 vector, created by a pooling layer.

LeNet’s dense block has three fully-connected layers, with 120, 84, and 10 outputs, respectively. Because we are still performing classification, the 10-dimensional output layer corresponds to the number of possible output classes. Implementing LeNet models with modern deep learning frameworks is remarkably simple.

## 4. Implementation and Results

### 4.1 ABIDE dataset

The rs-fMRI data of ASD and healthy subjects are downloaded from a large multisite data repository ABIDE (Autism Brain Imaging Data Exchange) (Http://Fcon_1000.Projects.Nitrc.Org/Indi/Abide/). The Autism Brain Imaging Data Exchange I (ABIDE I) is a multisite platform gathered from 17 international laboratories, shared some collected resting state functional magnetic resonance imaging (rs-fMRI), anatomical and phenotypic datasets. This dataset includes 1112 patients, from 539 individuals with ASD and 573 from typical controls (ages 7-64 years, median 14.7 years across groups). Till now, this data is used in many researches. The publications have shown its utility for capturing whole brain and regional properties of the brain connectome in Autism Spectrum Disorder (ASD). All data have been anonymized.

Accordingly, ABIDE II was established to further promote discovery science on the brain connectome in ASD. To date, ABIDE II involves 19 sites, overall donating 1114 datasets from 521 individuals with ASD and 593 controls (age range: 5-64 years). All datasets are anonymous, with no protected health information included.

There is not ASD/healthy label for some individuals present in ABIDE database. After removing these cases, 871 individuals of ABIDE I and 910 individuals of ABIDE II would be remained, for investigation in this study (X. Yang et al., 2019).

### 4.2 Preprocessing and Connectivity Matrix

The rs-fMRI data are slice time corrected, motion corrected, registered and normalized, using FSL software. The steps of preprocessing done for ABIDE I and ABIDE II databases are: 1) AC-PC Realignment, 2) Gray Matter, and White Matter Tissue Segmentation, 3) Non-linear registration to MNI152 space, 4) Normalization, 5) Resampling, 6) modulation, and 7) Smoothing with FWMH=4mm. For the task of brain parcellation, the ICA method is used (F. de Martino et al., 2007; Joel et al., 2011; Smith et al., 2009; Tohka et al., 2008). In other words, instead of obtaining average of the time series (BOLD signal) of some pre-defined regions, spatial maps output from ICA with the specific functional and anatomical interpretation (the locations of brain tissue acting synchronously and with the same activity pattern) are taken into account. ICA is a data-driven model which uses no a priori information about the brain and has been a popular approach in the analysis of fMRI data (Salimi-Khorshidi et al., 2014). In this study ICA decomposed the whole BOLD fMRI data into 39 regions according to MNI ATLAS.

Afterward, the BOLD signal of these 39 ROI is considered to compute their connectivity measures, by the statistical measures such as Pearson correlation, partial correlation (Saad et al., 2009) and tangent correlation (Dadi et al., 2019). The size of connectivity matrix is 39*39, according to number of ROIs. The Pearson correlation coefficient ranges from 1 to −1. 1 indicates that two ROIs are highly correlated, −1 indicates that two ROIs are anti-correlated. This step is done using Nilearn toolbox developed by MIT university, and as well by BrainIAK toolbox (Kumar et al., 2020). Nilearn is a python toolbox for statistical learning on neuroimaging data. In this study, connectivity matrix is obtained via tangent correlation (Pedregosa et al., 2011). See Appendix A for more details. This method is less frequently used, but has solid mathematical foundations and a variety of groups have reported good decoding performances with this framework. Connectivity matrices built with tangent space parametrization give an improvement compared to full or partial correlations.

### 4.3 Classifying the Graph Embedding Vectors

According to the abovementioned embedding features, we used three scenarios to check whether ASD detection can be improved by graph embedding algorithms or not. In first scenario, features are embedded vectors of the connectivity matrix using each of Node2Vec, Struc2Vec, and AWE method. Accordingly, a deep network with one channel input is used in this scenario. This channel input is an *d*×*N* matrix including the *d*-dimensional embedded vectors of all *N* = 39 nodes. The embedded vectors obtained by these methods has a dimension of *d* = 25, 64, and 128 respectively. But, in the second scenario, to take different properties of Node2vec algorithm, with different *p* and *q* values (*p* = 1, *q* = 1, *p* = 1, *q* = 4, and *p* = 4, *q* = 1), into account, a 3-channel deep network is applied. In the third scenario, after applying PCA on the result of Node2Vec algorithm, two matrices of the Graph2Img algorithm are considered as input of a 2-channel CNN. These three scenarios are schematically shown in Figure 7.

**Figure 7.**
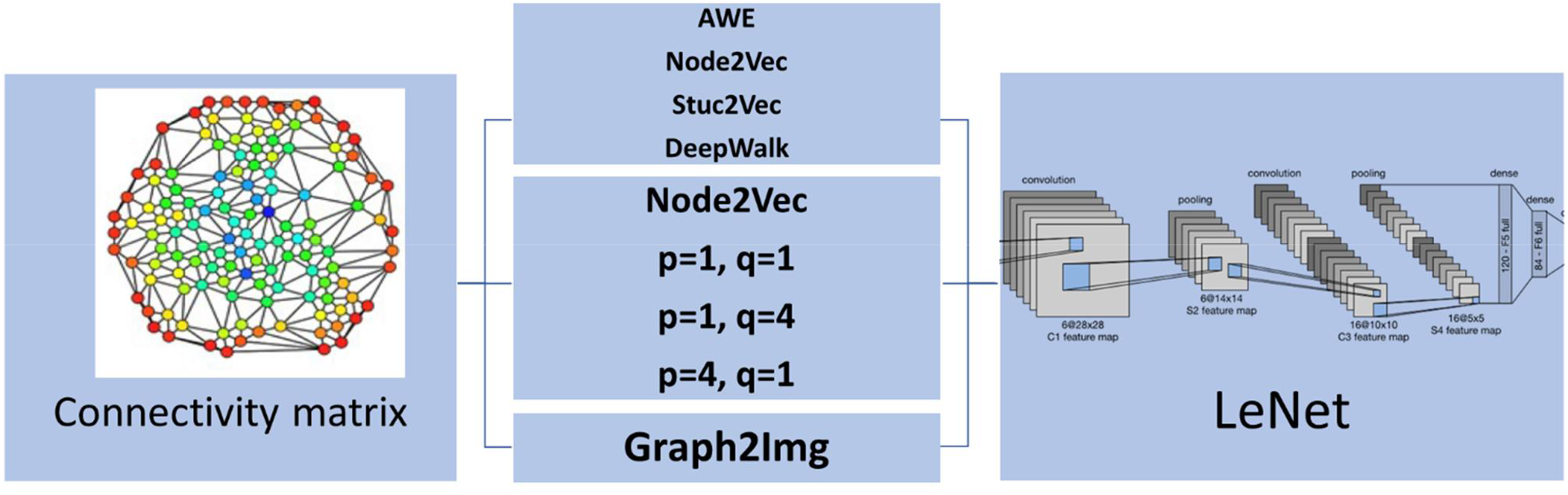
Three Scenarios of applying embedding vectors to detect ASD via LeNet

Indeed, at first, we tried to do the classification job through traditional kernel-base classifiers, like Support Vector Machine (SVM), but satisfactory results could not be obtained. The classifier could not show an accuracy better than chance. The advantage of CNN is that it is composed of an automatic feature extractor, that again extract features from the embedded vectors and, thus is a trainable classifier.

In all three scenarios, we customize the LeNet structure for our problem: In the 1^st^ scenario, there are one channel in the input layer, and the size of the embedded vector in each of Node2vec, Struct2vec, and AWE method determines dimensional of the input. These sizes are respectively equal to 25, 64, and 128.

Thus, in the 1^st^ scenario, the input layer of LeNet is a 39 * *d,d* = 25,64,128 image. In the 2^nd^ scenario the network has three channels. Each channel of the deep network consists of 39 * *d* neurons. In the third scenario, there are two channels, each consisting 10*10 neurons (*r* = 10). Lastly, in all three scenarios, there are two output neurons indicating healthy and autism brain.

The default LeNet network was modified according to the abovementioned dimensions of input/output. Furthermore, a dropout layer is employed for regularization at every hidden layer [33] with 0.8 keeping regularity. Another difference is the activation functions we used in LeNet to be ReLU functions, except for the ultimate layer, which uses a softmax function in such a way that a probability distribution over classes would be obtained. For the convolution-pooling block, we employ 64 filters at the first level, and as the signal is halved through the (2,2) max pooling layer, the number of filters in the subsequent convolutional layer is increased to 96 to compensate for the loss in resolution (Tixier et al., 2019). The number of trainable weights in this deep neural network doubles or triples in the 3^rd^ and 2^nd^ scenario.

The illustrated networks are used as the healthy/ASD classifier. Classification results would be reported in section 5 to compare them with previous researches in which deep network is used to classify the raw connectivity matrices.

### 4.4 Evaluation

To check the performance of our proposed ASD classifier working based on graph embedding techniques and deep machine learning methods, two kinds of cross-validation techniques are used. Indeed, these two techniques depend on how we choose training and test datasets. According to the properties of ABIDE database that consists of different sites we can do three different partitioning jobs: 1) dividing data of each site into *N* folds, and report accuracy of classification in individual sites, 2) Leave-One-Site-Out validation (distinctly for ABIDE I and II), 3) dividing all data of ABIDE I and II into *N* folds to report a typical *N*-fold cross validation. In all three approaches, the classification performance is assessed by accuracy, F-score, recall and precision. To report the accuracy of all data, statistically more reliable, 2^nd^ approach, i.e., Leave-One-Site-Out validation, is the most appropriate one. However, in this paper, the validation type (2) and (3) are considered in the report of the results.

Considering ASD detection as the goal of classifier, True Positive (TP) is defined as the percent of ASD subjects correctly classified as ASD. As well, the percent of ASD subjects, classified as healthy is referred to as False Negative (FN). Similarly, False Positive (FP) is the percent of healthy subjects decided to be ASD. Accordingly, the F-score measure is defined as follows:

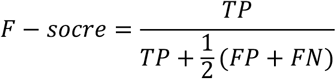

It is important for the classifier to detect all ASD subjects, so 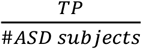 is referred to as recall.

Also, it is expected a classifier to have trusted positive detection, or in other word to be precise. Thus, precision is defined as 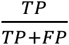. Because the size of subjects of two classes are not necessarily balanced, precision is a better measure of performance. Accordingly, another definition of F-score is based on recall, and precision:

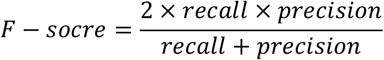

At last, to check how many subjects are correctly labelled, accuracy is a well-known measure.

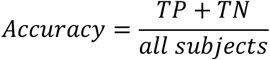

On the other hand, the time cost of training the classifier is another measure of the method under evaluation.

## 5. Results

Results of three scenarios for ABIDEI and ABIDE II database are presented in Table 1, using LeNet classifier. In the results of Table 1 validation of type 3 is considered: all subjects of each database are taken into account, then 5-fold and 10-fold cross-validation is applied. The average accuracy of these folds is reported for each scenario. Scenario 2 achieved the best performance in which a mean classification accuracy of 64% (recall 0.77%, precision 0.73%), and 66% (recall 80%, precision 80%) is obtained for ABIDE I and ABIDE II, respectively (in 10-fold cross-validation). The range of accuracy values were between 52% and 69% in individual folds. Based on the literature, this is not better than (Sherkatghanad et al., 2020) in which 70.22% accuracy is reported.

**Table 1.**
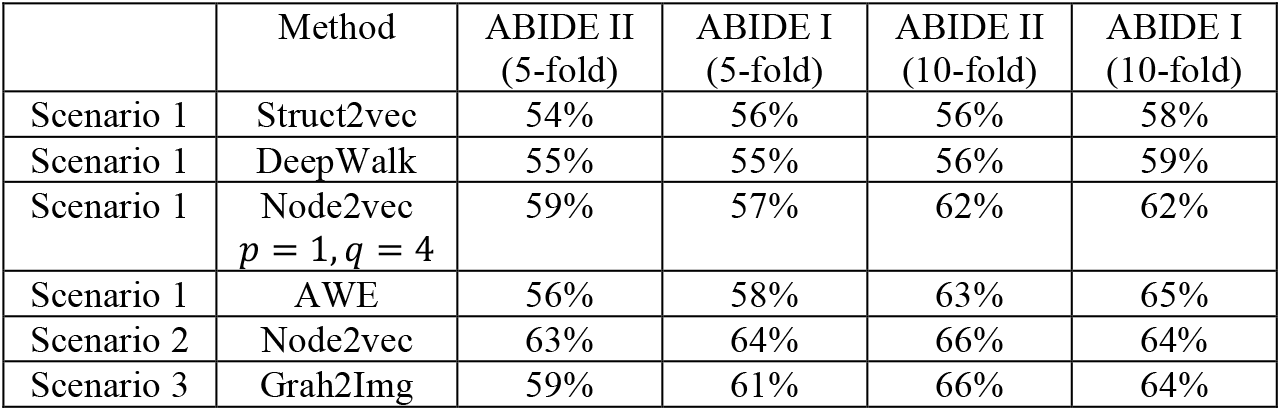
5 and 10-fold cross-validation results using different embedding methods and CNN classifier (LeNet)

The results of Table 1 show that the type of embedded features is effective in classification. But, as mentioned before, not given here, the results of SVM using embedded features is not better than those of (Sherkatghanad et al., 2020), in which raw connectivity matrix has been used in classification. In other words, it seems that it is the art of deep network classifier in reaching a good separation between ASD and healthy subjects, not the embedding features. So, the question is that whether feature embedding method was effective in ASD/healthy discrimination or not.

On the other hand, results of the leave-one-site-out cross validation is reported in Tables 2 to 5, respectively for the scenario 1 using AWE, scenario 2 (node2vec), and scenario 3 (Graph2Img). In this validation type, just AWE of scenario 1 is applied, due to its better performance in the k-fold cross validation procedure, against other embedding techniques. For each site, the LeNet CNN classifier is trained by data of other sites in each database, and has been tested on data of that site. Results of the ABIDEI and ABIDEII are distinctly presented. Number of the subjects of each site, number of the ASD subjects, accuracy and F-score of the proposed techniques, as well as those of (Heinsfeld et al., 2018; Sherkatghanad et al., 2020) (just for ABIDE I) are reported in the tables.

**Table 2.**
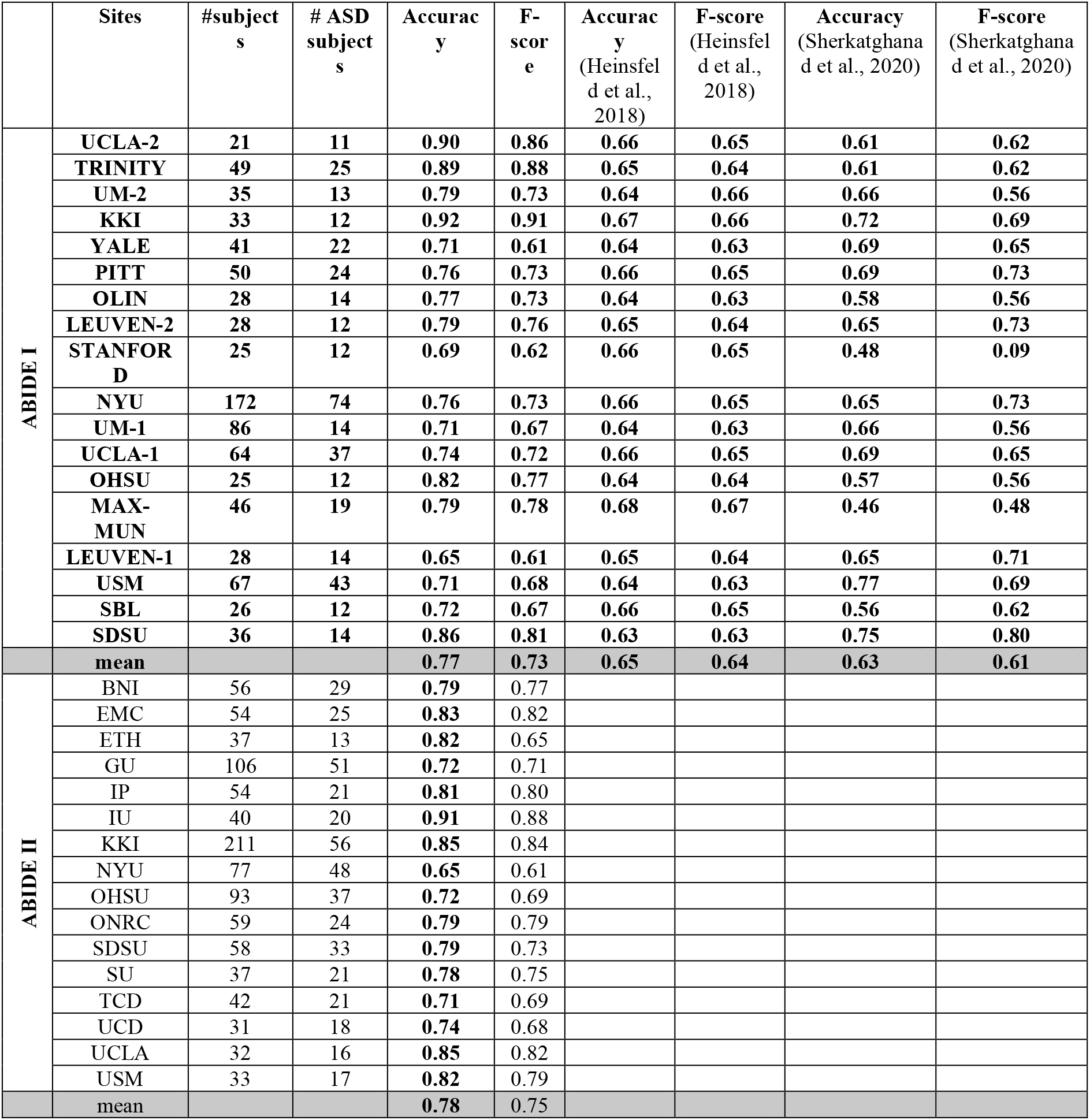
Leave-site-out cross-validation results using scenario 1 just by AWE and CNN classifier

**Table 3.** Leave-site-out cross-validation results using scenario 2 (Node2vec) and CNN classifier

|  | Sites | #subjects | # ASD subjects | Accuracy | F-score | Accuracy (Heinsfeld et al., 2018) | F-score (Heinsfeld et al., 2018) | Accuracy (Sherkatghana et al., 2020) | F-score (Sherkatghana et al., 2020) |
| --- | --- | --- | --- | --- | --- | --- | --- | --- | --- |
| **ABIDE I** | **UCLA-2** | **21** | **11** | **0.76** | **0.68** | **0.66** | **0.65** | **0.61** | **0.62** |
|  | **TRINITY** | **44** | **25** | **0.72** | **0.64** | **0.65** | **0.64** | **0.61** | **0.62** |
|  | **UM-2** | **34** | **13** | **0.71** | **0.63** | **0.64** | **0.66** | **0.66** | **0.56** |
|  | **KKI** | **33** | **12** | **0.66** | **0.64** | **0.67** | **0.66** | **0.72** | **0.69** |
|  | **YALE** | **41** | **22** | **0.81** | **0.76** | **0.64** | **0.63** | **0.69** | **0.65** |
|  | PITT | 50 | 24 | 0.76 | 0.74 | 0.66 | 0.65 | 0.69 | 0.73 |
|  | OLIN | 28 | 14 | 0.87 | 0.83 | 0.64 | 0.63 | 0.58 | 0.56 |
|  | LEUVEN-2 | 28 | 12 | 0.78 | 0.70 | 0.65 | 0.64 | 0.65 | 0.73 |
|  | STANFORD | 25 | 12 | 0.82 | 0.82 | 0.66 | 0.65 | 0.48 | 0.09 |
|  | NYU | 172 | 74 | 0.63 | 0.58 | 0.66 | 0.65 | 0.65 | 0.73 |
|  | UM-1 | 86 | 14 | 0.67 | 0.63 | 0.64 | 0.63 | 0.66 | 0.56 |
|  | UCLA-1 | 64 | 37 | 0.65 | 0.54 | 0.66 | 0.65 | 0.69 | 0.65 |
|  | OHSU | 25 | 12 | 0.77 | 0.72 | 0.64 | 0.64 | 0.57 | 0.56 |
|  | MAX-MUN | 46 | 19 | 0.60 | 0.53 | 0.68 | 0.67 | 0.46 | 0.48 |
|  | LEUVEN-1 | 28 | 14 | 0.85 | 0.82 | 0.65 | 0.64 | 0.65 | 0.71 |
|  | USM | 67 | 43 | 0.76 | 0.69 | 0.64 | 0.63 | 0.77 | 0.69 |
|  | SBL | 26 | 12 | 0.76 | 0.70 | 0.66 | 0.65 | 0.56 | 0.62 |
|  | SDSU | 27 | 14 | 0.73 | 0.65 | 0.63 | 0.63 | 0.75 | 0.80 |
|  | mean |  |  | 0.73 | 0.68 | 0.65 | 0.64 | 0.63 | 0.61 |
| ABIDE II | BNI | 56 | 29 | 0.66 | 0.60 |  |  |  |  |
|  | EMC | 54 | 25 | 0.80 | 0.77 |  |  |  |  |
|  | ETH | 34 | 13 | 0.74 | 0.65 |  |  |  |  |
|  | GU | 106 | 51 | 0.64 | 0.59 |  |  |  |  |
|  | IP | 54 | 21 | 0.74 | 0.69 |  |  |  |  |
|  | IU | 34 | 20 | 0.75 | 0.67 |  |  |  |  |
|  | KKI | 34 | 56 | 0.79 | 0.70 |  |  |  |  |
|  | NYU | 77 | 48 | 0.68 | 0.61 |  |  |  |  |
|  | OHSU | 93 | 37 | 0.63 | 0.56 |  |  |  |  |
|  | ONRC | 49 | 24 | 0.80 | 0.75 |  |  |  |  |
|  | SDSU | 58 | 33 | 0.67 | 0.58 |  |  |  |  |
|  | SU | 54 | 21 | 0.70 | 0.64 |  |  |  |  |
|  | TCD | 39 | 21 | 0.76 | 0.74 |  |  |  |  |
|  | UCD | 31 | 18 | 0.77 | 0.72 |  |  |  |  |
|  | UCLA | 32 | 16 | 0.80 | 0.76 |  |  |  |  |
|  | USM | 33 | 17 | 0.70 | 0.64 |  |  |  |  |
|  | mean |  |  | 0.72 | 0.66 |  |  |  |  |

**Table 4.**
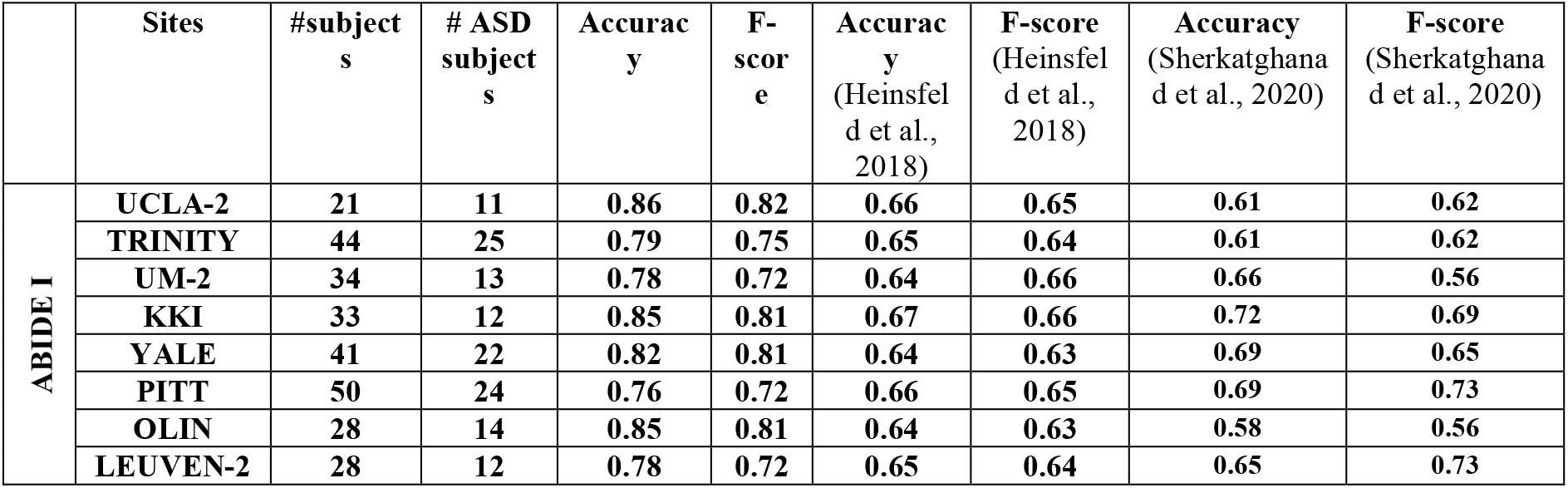
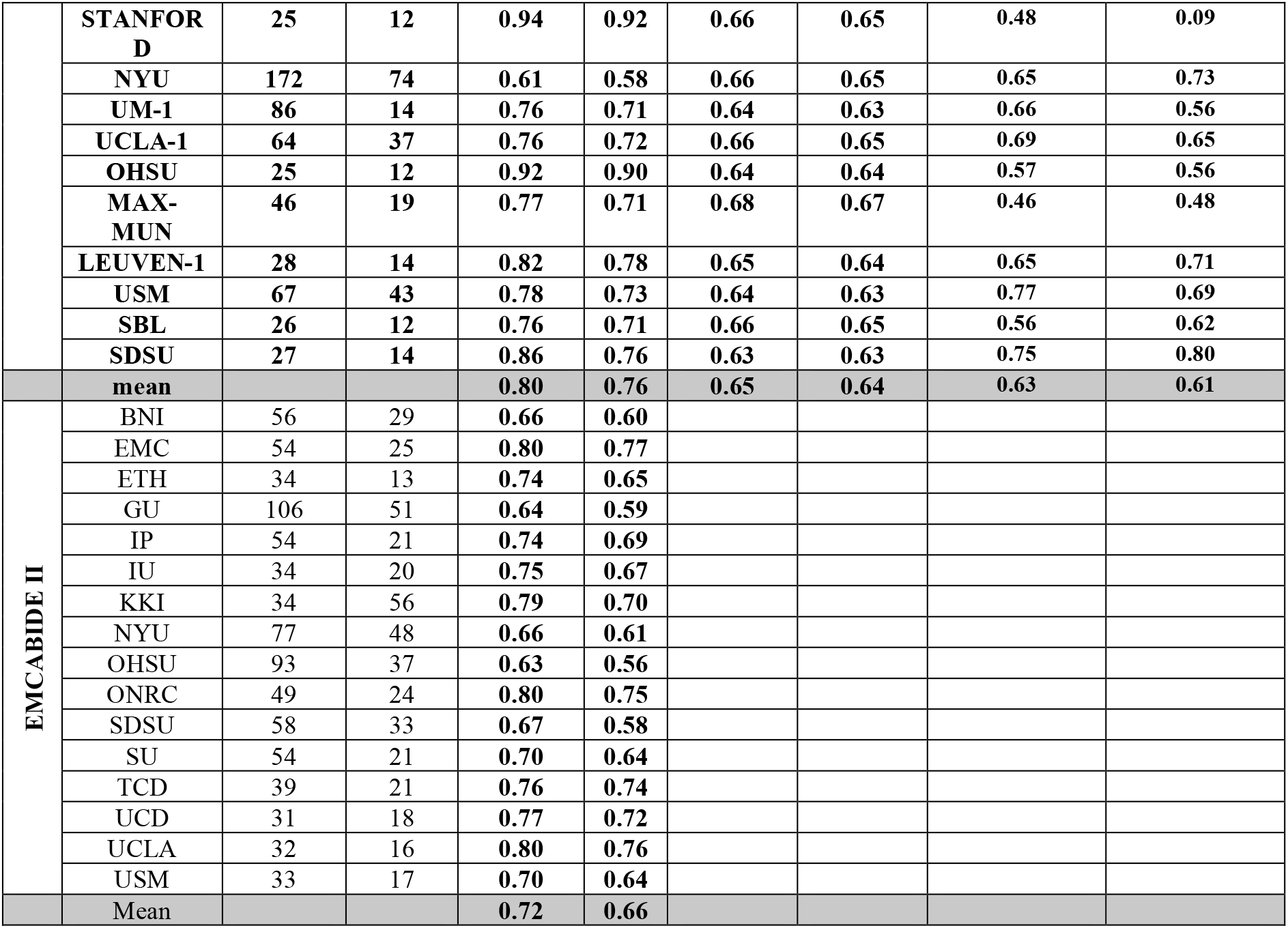
Leave-site-out cross-validation results using scenario 3 (Graph2Img) and CNN classifier

**Table 5.** Summary of best performance values and computational time for ABIDE I, in compared to literature.

|  | Mean Accuracy<br>(10-fold cross-validation) | Mean Accuracy<br>(Leave-one-site-out) | Computation Time |
| --- | --- | --- | --- |
| Heinsfeld<br>(Heinsfeld et al., 2018) | 70% | 65% | Over 32h |
| SherkatGhanad<br>(Sherkatghanad et al., 2020) | 70.22% | 63% | 12h 30m |
| Proposed method | 66% | 80% | 2h 47m 20s |

As shown in Figure 8, these results in compared to (Heinsfeld et al., 2018; Sherkatghanad et al., 2020), that achieved the best results in the literature so far, depicts that the embedding-based CNN can outperform the other supervised methods. From this point of view, these results are in favor of embedding features, not just the Deep Network.

**Figure 8.**
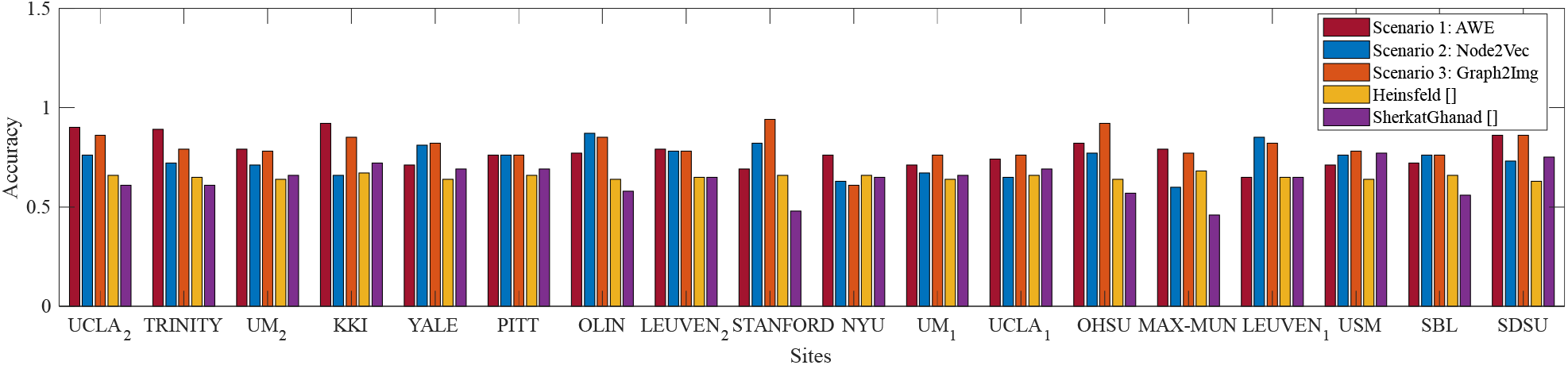
Box plot of leave-out-site accuracy to compare different embedding scenarios and (Heinsfeld et al., 2018; Sherkatghanad et al., 2020) vs. sites, for ABIDE I dataset.

**Figure 9.**
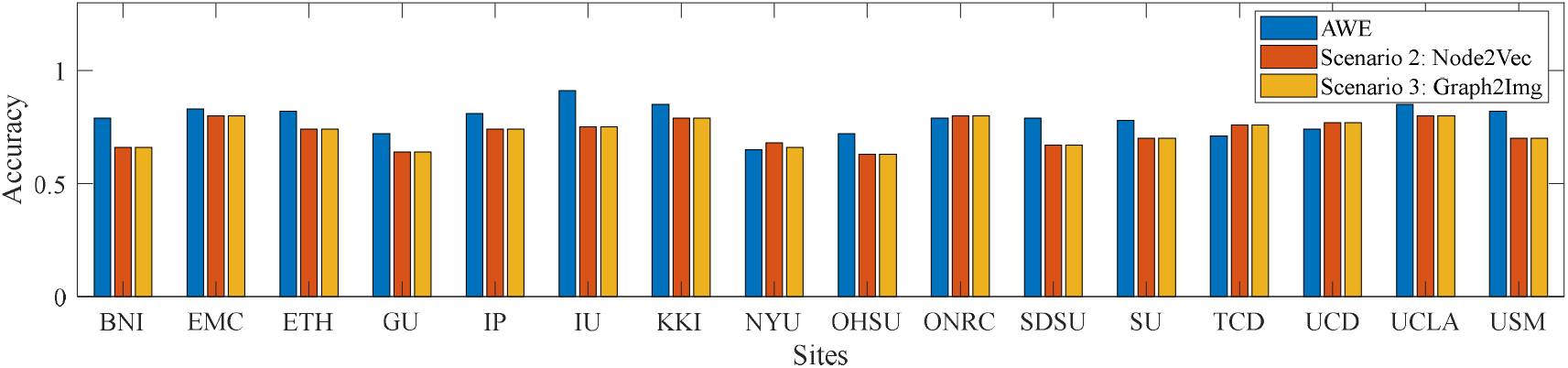
Box plot of leave-out-site accuracy to compare different embedding scenarios vs. sites, for ABIDE II dataset.

Therefore, some points worth considering in the results of these two validation methods:

1. Embedding features could not improve the results of the k-fold cross validation, but is able to improve the results of leave-one-site-out one.
2. The accuracy of CNN in classifying ASD subjects of each site is different when using graph embedding methods. In average, all embedding scenarios could improve the results, in comparison to using the raw connectivity matrices, in the leave-site-out validation manner (Heinsfeld et al., 2018; Sherkatghanad et al., 2020). The best embedding technique seems to be Graph2Img that increase the 65% (Heinsfeld et al., 2018) and 63% (Sherkatghanad et al., 2020) results to 80%.
3. Each graph embedding scenario has significantly improved the results of some sites, but not the all of sites. The AWE technique is not successful about the fMRI data of University of Utah School of Medicine (USM), for which (Sherkatghanad et al., 2020) acts well. As well (Sherkatghanad et al., 2020) and/or (Heinsfeld et al., 2018) outperforms the embedding scenario 2 (three channel node2vec with three values of p and q) in Kennedy Krieger Institute, Baltimore (KKI) data, New York University Langone Medical Center (NYU), Ludwig Maximilian University Munich (MAX-MUN), USM, and San Diego State University (SDSU) data. Even, for the embedding scenario 3 (i.e., Graph2Img) there is a site for which the accuracy of ASD classification is lower than (Sherkatghanad et al., 2020).
4. For the ABIDE II database, scenario 1 (AWE method) reached the best mean accuracy. The best individual site result, also is dedicated to AWE method for the KKI database.

However, the most dominant advantage of the proposed algorithm is its training time. Using a system with two Intel Xeon E5-2620 processors with 24 cores running at 2 GHz and 48 GB of RAM. As well, 1 Tesla K40 GPU with 2880 CUDA cores and 12 GB of RAM was used to accelerate training.

In such a way, the entire training time took about 200 minutes. In Table 5 the training time of (Heinsfeld et al., 2018; Sherkatghanad et al., 2020) and our proposed method is compared. This achievement is due to the dimension reduction property of the graph embedding methods, decreasing the dimensionality of the CNN input.

The results show that the proposed algorithm, using embedded vectors of connectivity graph, and the CNN classifier, outperforms the previous studies in identification of autism spectrum disorder, from both speed and accuracy point of view.

## 6. Conclusion and Discussion

Since the functioning of the brain is accompanied by interactions and connection between different functional areas, discrimination of healthy and autism behaviors could be done by assessment of the brain network dynamics (Kim et al., 2017). Indeed, cognitive disorders emerges because of the alteration of dynamic relationships between pairs of specific brain regions. However, we claim that a powerful learning method considering the coupling, similarity, or causality and synchronizing intensity between specific brain regions could be able to detect cognitive impairment.

For a complete comparison, we considered the literature where functional connectivity matrix is used to discriminant healthy and autistic subjects, based on ABIDE database, either using conventional classifiers, or deep networks, or even statistical tests. The result of these comparisons is shown in Figure 8, illustrating that embedded features achieved better results than other feature extraction methods, and, as well, deep neural networks hold much greater promises than conventional classifiers. In other words, since AWE (Scenario 1), Graph2Img (Scenario 3) and Multi-parameter Node2Vec (Scenario 2) algorithms gain better classification results with CNN classifier (in leave-one-site-out validation), we claim that embedded features involving structure of the functional connectivity of brain could be more convenient in ASD detection. The classification results (in k-fold cross validation), although not high (66% and 64% for ABIDEI and II) enough to be appropriate for clinical usage, show that there are strong alterations in brain connections during autism disorder.

Our 10-fold cross-validation best average result is 65% in compared to (Heinsfeld et al., 2018) that is about 65%. Instead, as shown in Tables 2 to 4, and Figure 8, for leave-one-site-out, both the mean accuracy over sites, and the most of individual accuracy of sites, our proposed method is clearly much better than (Heinsfeld et al., 2018) and (Sherkatghanad et al., 2020). These results show that, as the sample size decreases (5-fold, 10-fold cross-validation results, and leave-one-site-out), the gap between performance of the embedding vectors, and the raw connectivity matrix increase. This implies that using embedding vectors is an effective idea, but still needs more investigation to find the more suitable graph representation method. The reason is clearly the intrinsic complexity of brain function.

There are two messages in the obtained results: First, the intrinsic phenotypical properties of subjects within each site, lead to a specific structure in their connectivity graph, in addition to the distinct indicator of ASD/healthy. Different embedding techniques acquire some of these properties. Second, a suitable combination of graph embedding techniques is the alternative approach to take all graph similarities in the ASD group regardless the phenotypes.

The better mean accuracy of leave-one-site-out validation technique compared to that of k-fold cross validation again tells us about the variance of the graph structures between sites due to the within site phenotypes. In such a way, in a random group, finding the common structures just relevant to ASD would be too difficult for an embedding technique. It is the main reason prevents the embedding techniques capture a better result than the raw connectivity matrices.

In this paper, we showed that using structural graph representation algorithms, it is possible to classify subject groups based on connectivity fingerprints of brain regions. However, we did not analyze the results to obtain knowledge about these alterations were of what kind, and where they occur. This is drawback of our proposed algorithm that we could not identify ROIs that alter connectivity strength values. In fact, the main point of a suitable embedding algorithm for brain network is that the representations they emerge would be neurobiologically plausible and meaningful. From this point of view, we can predict the mechanism and causes underlying an impaired brain network during mental disorders. This is our future concern.

## 7. Appendix: Computing the covariance tangent space

Nilearn Python library provides the tangent-space parametrization of covariance matrices. In this appendix it is described how this parametrization is computed, with required formulation (Dadi et al., 2019). The algorithm is made of two steps: First covariance matrix of BOLD signal of each subject is computed, and the group average matrix over all subjects are obtained. Second, this average matrix is used to transform covariance matrix to a space where ensures a well-conditioned connectivity matrix.

To estimate the average covariance matrix, the Ledoit and Wolf (Ledoit & Wolf, 2004) estimator is a good choice (Brier et al., 2015; Varoquaux & Craddock, 2013). This namely Frechet mean is calculated according to the geometry of covariance matrices (Pennec et al., 2006; Varoquaux et al., 2010), in order to minimize a cost function. This cost function is depicted in algorithm 3 of Fletcher and Joshi (Fletcher & Joshi, 2007). But, Euclidean mean with the following formula give almost the same performance.

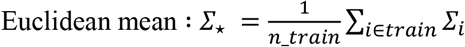

After computing this average matrix *Σ*_*_, covariance matrix should be transformed in the tangent-space representation. This is done by first normalizing the covariance matrix of each subject, and then mapping it to a suitable space.

1. After eigenvalue decomposing of *Σ*_*_ = *U^T^*Δ*U*, where *U* contains eigenvectors and Δ is a diagonal matrix containing eigenvalues, we can compute the normalized matrix of each BOLD signal via

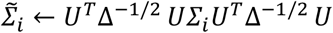
2. After eigenvalue decomposing of 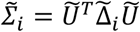, we can compute the matrix logarithm 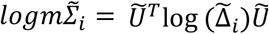, where the logarithm is applied to the diagonal elements of 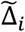. Finally, the 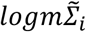 is the resulting matrix, and can be considered as the tangent connectivity matrix.

